# SGK phosphorylates Cdc25 and Myt1 to trigger cyclin B-Cdk1 activation at the meiotic G2/M transition

**DOI:** 10.1101/499640

**Authors:** Daisaku Hiraoka, Enako Hosoda, Kazuyoshi Chiba, Takeo Kishimoto

## Abstract

The kinase cyclin B-Cdk1 complex is a master regulator of M-phase in both mitosis and meiosis. At the G2/M transition, cyclin B-Cdk1 activation is initiated by a trigger that reverses the balance of activities between Cdc25 and Wee1/Myt1, and is further accelerated by autoregulatory loops. In somatic cell mitosis, this trigger was recently proposed to be the cyclin A-Cdk1/Plk1 axis. However, in the oocyte meiotic G2/M transition, in which hormonal stimuli induce cyclin B-Cdk1 activation, cyclin A-Cdk1 is non-essential and hence the trigger remains elusive. Here, we show that SGK directly phosphorylates Cdc25 and Myt1 to trigger cyclin B-Cdk1 activation in starfish oocytes. After hormonal stimulation of the meiotic G2/M transition, SGK is activated by cooperation between the Gβγ-PI3K pathway and an unidentified pathway downstream of Gβγ, called the atypical Gβγ pathway. These findings identify the trigger in oocyte meiosis and provide insights into the role and activation of SGK.

## Introduction

Activation of cyclin-dependent kinase 1 complexed with cyclin B (cyclin B-Cdk1) induces entry into M-phase during somatic cell mitosis and germ cell meiosis^1–3^. Cyclin B-Cdk1 is regulated by synthesis and degradation of cyclin B, and by inhibitory phosphorylation of Cdk1 at Thr14 and Tyr15. These residues are phosphorylated by kinases belonging to the Wee1/membrane-associated tyrosine/threonine 1 (Myt1) family, but dephosphorylated by cell division cycle 25 (Cdc25)^4^. Cyclin B accumulates before and/or during M-phase entry. However, Wee1/Myt1 activity is dominant over Cdc25 activity before M-phase; therefore, cyclin B-Cdk1 remains inactive due to inhibitory phosphorylation. At M-phase entry, a small population of cyclin B-Cdk1 is first activated by a trigger that reverses the balance between Cdc25 and Wee1/Myt1 activities, thereby making Cdc25 activity predominant. Thereafter, cyclin B-Cdk1 itself further accelerates dephosphorylation of Cdk1 via feedback loops, leading to maximal activation^4–8^. Although the molecular identity of the trigger of cyclin B-Cdk1 activation has received great attention, it remains elusive.

In mitotic M-phase entry (G2/M transition), the trigger may be affected by various factors, such as checkpoints^5,9,10^, and may involve redundant or stochastic processes^5,9^. At least in normal mitotic cell cycles, polo-like kinase 1 (Plk1) is activated by Aurora A in a cyclin A-Cdk1-dependent manner, and in turn phosphorylates Cdc25 to trigger activation of cyclin B-Cdk1^11–13^. Consistently, the cyclin A-Cdk1/Plk1 axis functions as the trigger for cyclin B-Cdk1 activation in the first embryonic cell division cycle^14^.

In contrast with mitotic cell cycles in proliferating somatic cells, meiotic cell cycles in animal oocytes mostly arrest at prophase of meiosis I (prophase-I), which corresponds to late mitotic G2 phase^15^. Release from this arrest is also induced by cyclin B-Cdk1, which is activated downstream of extracellular hormonal stimuli^16^; hence this is known as the meiotic G2/M transition. However, this transition has not been reported to require cyclin A^16–21^. Furthermore, Plk1^14,22,23^ and Aurora^24–26^ are non-essential in some oocytes. Thus, mechanisms other than the cyclin A-Cdk1/Plk1 axis likely trigger cyclin B-Cdk1 activation at the meiotic G2/M transition in oocytes.

In vertebrate oocytes, meiotic G2 arrest requires cAMP-dependent protein kinase A (PKA), and downregulation of this kinase leads to meiotic G2/M transition^16,20,27,28^. In mice, PKA appears to directly upregulate Wee1B and downregulate Cdc25; however, it remains unclear how downregulation of PKA reverses the balance between Wee1B and Cdc25 activities to trigger cyclin B-Cdk1 activation^29^. In *Xenopus*, the hormone progesterone stimulates protein synthesis of Moloney sarcoma oncogene (Mos) and cyclin B, which redundantly trigger cyclin B-Cdk1 activation^30^. This redundant process is possibly mediated in two manners: activating phosphorylation of Cdc25 by Mos/mitogen activated protein kinase (MAPK) cascade, and inhibitory phosphorylation of Myt1 by newly synthesized cyclin B-associated Cdk1^23^. However, the molecular link between downregulation of PKA and *de novo* protein synthesis of Mos and cyclin B remains elusive^31,32^.

In contrast with vertebrate oocytes, the mechanisms by which activation of cyclin B-Cdk1 is triggered at the meiotic G2/M transition have been well studied in invertebrate starfish oocytes since the existence of a trigger kinase was first reported^16,21,33^. In starfish, cAMP and PKA are likely not involved in meiotic G2 arrest^34^. The physiological maturation-inducing hormone 1-methyladenine (1-MeAde) induces the meiotic G2/M transition^35^ without a requirement for new protein synthesis^36^. Cyclin A, Wee1, and Mos are not present, and Aurora and Plk are not required for cyclin B-Cdk1 activation, in the meiotic G2/M transition^14,19,25,37^. In unstimulated oocytes, cyclin B is already accumulated^38^, but Myt1 inactivates cyclin B-Cdk1^39^. Stimulation by 1-MeAde is mediated by an unidentified cell surface G-protein-coupled receptor (GPCR), which induces dissociation of Gβγ from Gαi on the plasma membrane^40–43^. Gβγ binds to and activates phosphoinositide-3-kinase (PI3K), resulting in production of phosphatidylinositol 3,4,5-triphosphate (PI-345P_3_)^44,45^. Dependent on PI-345P_3_, phosphoinositide-dependent kinase 1 (PDK1) and target of rapamycin complex 2 (TORC2) phosphorylate Akt in its activation loop (A-loop) and hydrophobic motif (HM), respectively, leading to its activation^46,47^. Thereafter, Akt inactivates Myt1 by directly phosphorylating Ser75 (ref. 39) and activates Cdc25 by directly phosphorylating multiple residues including Ser188 (ref. 48), resulting in reversal of the balance of Cdc25 and Myt1 activities, and consequently initial activation of cyclin B-Cdk1 (refs 33,39). These observations suggest that Akt likely represents the trigger kinase at the meiotic G2/M transition in starfish oocytes^39^.

Nevertheless, our recent observations suggested that the PI3K-Akt pathway is insufficient^48^. Expression of a constitutively active form of Akt (CA-Akt) induces the meiotic G2/M transition, but the required expression level of CA-Akt is 40-fold higher than that of endogenous Akt. Furthermore, expression of a constitutively active form of PI3K (CA-PI3K) induces Akt activation, but results in low levels of Cdc25 and Myt1 phosphorylation and therefore fails to induce the meiotic G2/M transition. We reported that another pathway, called “the atypical pathway”, which is activated by Gβγ in parallel with PI3K activation, compensates for this insufficiency^48^. Although the molecular details of this pathway remain unclear, we proposed that it cooperates with the PI3K-Akt pathway to trigger cyclin B-Cdk1 activation^48^. One possibility is that another kinase, which can redundantly phosphorylate Akt substrates, functions in these pathways to trigger cyclin B-Cdk1 activation.

One such candidate kinase is serum- and glucocorticoid-regulated kinase (SGK)^49,50^. SGK is a member of the AGC kinase family, which includes Akt^51^, and has the same substrate consensus motif (RXRXXS/T, where X represents any amino acid) as Akt^52^. In mammalian cells, similar to Akt, SGK family members are activated in a PI3K-dependent manner, dependent on phosphorylation of the A-loop and HM by PDK1 and TORC2, respectively^52,53^. However, the contribution of SGK to the meiotic G2/M transition has not been investigated.

Here, we carefully evaluated whether Akt and SGK play redundant roles in the meiotic G2/M transition in starfish oocytes. We showed that SGK is necessary and sufficient for the initial phosphorylation of Cdc25 and Myt1 to trigger cyclin B-Cdk1 activation upon 1-MeAde stimulation. By contrast, although contradictory to our previous report^39^, inhibition of Akt did not affect the meiotic G2/M transition. Several experiments supported the importance of SGK, rather than of Akt. This discrepancy is discussed. Furthermore, as upstream pathways, we found that PI3K and the atypical Gβγ pathway cooperatively activate SGK. All these observations clearly explain the insufficiency of the PI3K-Akt pathway. Thus, our findings indicate that SGK is the trigger kinase at the meiotic G2/M transition in starfish oocytes, and provide new insights into SGK activation.

## Results

### A newly generated phospho-specific antibody detects A-loop phosphorylation of endogenous starfish Akt (sfAkt)

In previous studies^47,48^, we monitored phosphorylation of the HM by immunoblotting as a marker of starfish Akt (sfAkt) activity because attempts to generate an antibody that detects the phosphorylated A-loop were unsuccessful. Nevertheless, it is important to analyze A-loop phosphorylation to precisely evaluate Akt activation. Thus, we again attempted to generate a phospho-specific antibody against a 17-amino acid phospho-peptide derived from the A-loop of sfAkt that includes phospho-Thr at the site of phosphorylation by PDK1 (Fig. 1a). In immunoblot analysis of whole-oocyte samples using this antibody, phosphorylation of sfAkt was detectable following 1-MeAde stimulation only when Akt was overexpressed (Fig. 1b, closed arrowhead). When endogenous Akt was concentrated by immunoprecipitation using an antibody raised against a C-terminal fragment of sfAkt (anti-sfAkt-C antibody), the anti-phospho-A-loop antibody detected a band at the same size as Akt after 1-MeAde stimulation (Fig. 1c for input and flow-through, d for precipitate, closed arrowhead). The band disappeared upon treatment with the PDK1 inhibitor BX795 (Fig. 1d, closed arrowhead), indicating that the anti-phospho-A-loop antibody successfully detected A-loop phosphorylation of endogenous Akt by PDK1.

**Fig. 1.**
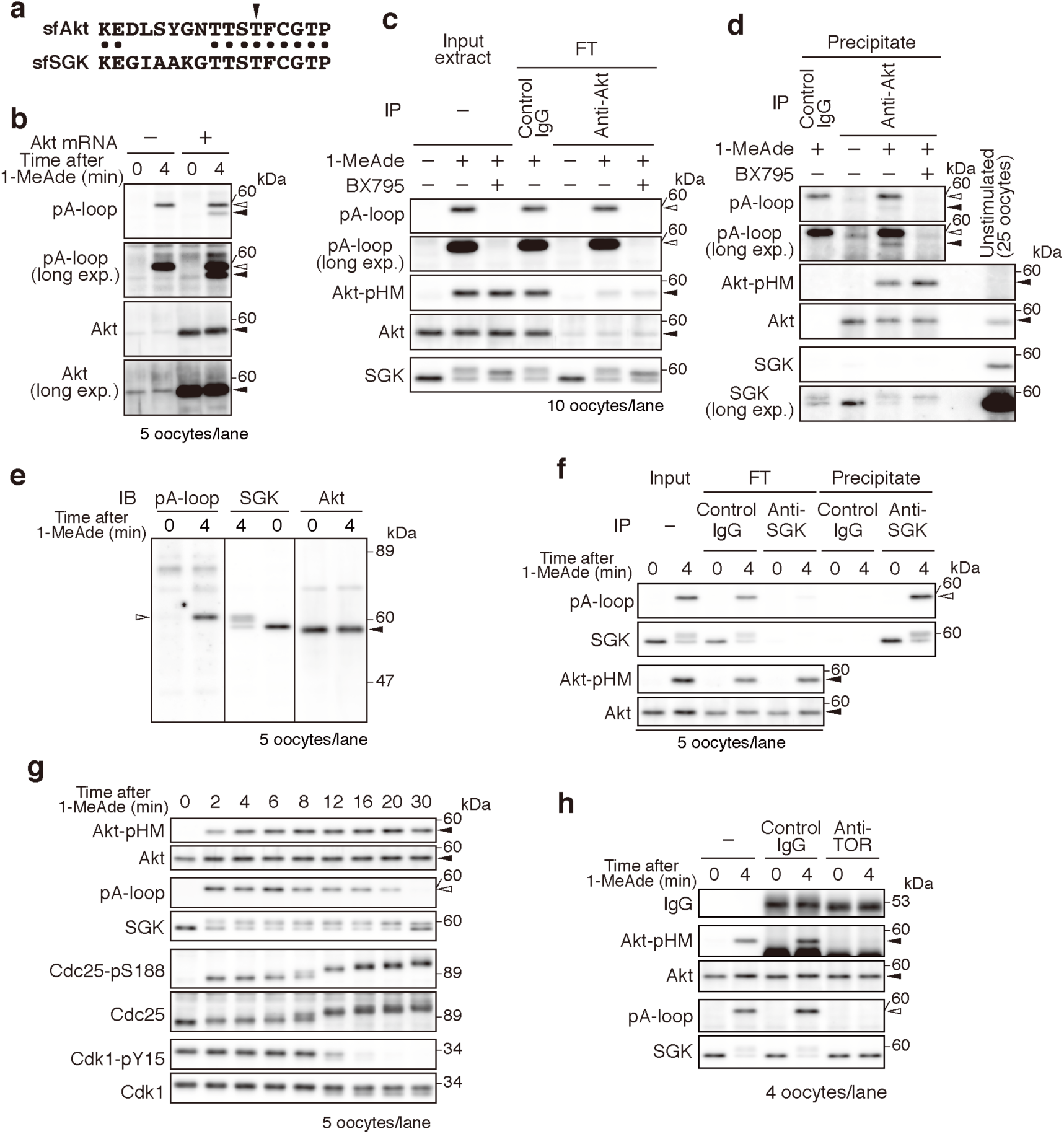
SGK is activated after 1-MeAde stimulation in a PDK1- and TORC2-dependent manner. **a** The region of sfAkt used to raise the anti-phospho-A-loop antibody (upper, Lys304–Pro320 of sfAkt) is aligned with the corresponding region of sfSGK (Lys301–Pro317). The arrowhead indicates the residue phosphorylated by PDK1 (Thr315 in sfAkt and Thr312 in sfSGK). Dots indicate identical residues. **b** A-loop phosphorylation of endogenous sfAkt is undetectable level in whole oocyte extracts. Uninjected oocytes and oocytes injected with mRNA encoding untagged sfAkt were treated with 1-MeAde, and analyzed by immunoblotting with the anti-phospho-A-loop (pA-loop) and anti-sfAkt-C (Akt) antibodies. Closed and open arrowheads indicate the positions of Akt and A-loop-phosphorylated SGK (see **e** and **f**), respectively. **c, d** A-loop phosphorylation of endogenous Akt is detectable in immunoprecipitates. Oocytes were incubated with 1-MeAde in the presence or absence of a PDK1 inhibitor, BX795, lysed, and subjected to immunoprecipitation with control IgG or the anti-sfAkt-C antibody. Input and flow-through (FT) extracts (**c**), and precipitates (**d**) were analyzed by immunoblotting (Akt-pHM represents phosphorylation of Ser477 in the HM of Akt). A-loop phosphorylation of non-specifically included SGK was detected in control and Akt immunoprecipitates (**d**, open arrowhead). **e** Unstimulated and 1-MeAde-treated oocytes were subjected to immunoblotting with anti-phospho-A-loop (left), anti-sfSGK-C (middle), and anti-sfAkt-C (right) antibodies. **f** The anti-phospho-A-loop antibody against sfAkt cross-reacts with A-loop-phosphorylated endogenous SGK. Unstimulated oocytes were treated with 1-MeAde. Immunoprecipitation was performed with control IgG or the anti-sfSGK-C antibody. Input, flow-through, and precipitates were analyzed by immunoblotting. **g** SGK is activated simultaneously with Akt activation. Oocytes were treated with 1-MeAde and collected at the indicated time points. Phosphorylation of Akt (Akt-pHM), SGK (pA-loop), Cdc25 (pS188), and Cdk1 (pY15) were analyzed by immunoblotting. GVBD occurred around 20 min. **h** SGK activation in starfish oocytes depends on TOR. Unstimulated oocytes were injected with control IgG or the anti-TOR antibody, treated with 1-MeAde, and analyzed by immunoblotting. The data are representative of two independent experiments in **b**, **e**, and **f**; and three independent experiments in **c**, **d**, and **g**. In all panels, closed and open arrowheads indicate the positions of Akt and A-loop-phosphorylated sfSGK, respectively.

### SGK is activated after 1-MeAde stimulation in a PDK1- and TORC2-dependent manner

The phospho-A-loop antibody detected another band that migrated slightly slower (~60 kDa) than Akt (Fig. 1b, c, open arrowhead). This band was still observed in flow-through samples in which Akt had been removed by immunoprecipitation (Fig. 1c, open arrowhead) and disappeared in the presence of BX795 (Fig. 1c, open arrowhead). Thus, we speculated that the band corresponds to PDK1-dependent A-loop phosphorylation of a kinase other than Akt.

In parallel with our research, Hosoda et al. found that starfish SGK (sfSGK) is activated after 1-MeAde stimulation in starfish oocytes in a study of intracellular pH regulation^54^. They isolated a cDNA encoding sfSGK and collaborated with us to generate an anti-sfSGK neutralizing antibody raised against a C-terminal peptide of sfSGK including the HM (anti-sfSGK-C antibody). A predicted molecular weight of sfSGK is 56 kDa, and eleven of the seventeen residues in the antigen peptide used to raise the anti-phospho-A-loop antibody against sfAkt are conserved in the A-loop of SGK (Fig. 1a). Thus, the other band detected by the anti-phospho-A-loop antibody may correspond to A-loop phosphorylation of SGK. The mobility of this band was the same as that of the top band of SGK detected by the anti-sfSGK-C antibody in 1-MeAde-treated oocytes (Fig. 1e, open arrowhead). In addition, the band disappeared after depletion of SGK from oocyte extracts (Fig. 1f), whereas it was detected in a sample of immunoprecipitated SGK (Fig. 1f), indicating that the anti-phospho-A-loop antibody also detects A-loop phosphorylation of sfSGK. Taken together, these results confirmed that SGK is activated after 1-MeAde stimulation.

SGK and Akt phosphorylate a Ser or Thr residue in the same consensus motif^52^ and may therefore play a redundant role in 1-MeAde signaling. We next analyzed the dynamics of SGK activation in starfish oocytes in detail. As previously reported^48^, phosphorylation of Akt in the HM and phosphorylation of Cdc25 at Ser188 were detected within 2 min after 1-MeAde addition (Fig. 1g). SGK was activated at a similar time (Fig. 1g, pA-loop). Thereafter, Tyr15 of Cdk1 was dephosphorylated, which is indicative of cyclin B-Cdk1 activation (Fig. 1g, 12 min and later). In mammalian cells, SGK is activated by phosphorylation of its A-loop and HM, which is catalyzed by PDK1 and mTORC2, respectively^52,53^. In addition, A-loop phosphorylation depends on HM phosphorylation^52^. In starfish oocytes, a mobility shift of SGK was still observed even when A-loop phosphorylation was abolished by BX795 (Fig. 1c). Using chemical inhibitors, Hosoda et al. suggested that this mobility shift represents TORC2-dependent phosphorylation^54^. Consistently, inhibition of TOR by microinjection of an anti-TOR neutralizing antibody^47^ completely abolished the mobility shift of SGK after 1-MeAde stimulation (Fig. 1h). In these oocytes, A-loop phosphorylation of SGK was also abolished (Fig. 1h). Therefore, A-loop phosphorylation of SGK likely depends on TORC2-dependent HM phosphorylation in starfish oocytes, as reported in mammalian cells.

### SGK, but not Akt, is required for the meiotic G2/M transition

Next, we examined the involvement of Akt and SGK in the meiotic G2/M transition. To this end, we specifically inhibited each kinase using antibodies. Non-specific bands were detected by immunoblotting of starfish oocytes with the anti-sfAkt-C antibody (Fig. 2a), but were not immunoprecipitated by this antibody (Fig. 2a). This indicates that the anti-sfAkt-C antibody does not interact with these non-specific proteins in non-denaturing conditions. In addition, although immunoprecipitates obtained using the anti-sfAkt-C antibody contained a trace amount of SGK (Fig. 1d, SGK with long exposure), this was not due to cross-reaction of SGK with this antibody because a similar amount of SGK was detected in a control precipitation obtained using a control IgG (Fig. 1d, SGK with long exposure). Furthermore, the amount of SGK was not reduced in extracts after immunoprecipitation of Akt (Fig. 1c). These observations confirm the specificity of the anti-sfAkt-C antibody. The anti-sfSGK-C antibody specifically detected SGK by immunoblotting (Fig. 1e). In addition, the protein level of Akt remained constant before and after depletion of SGK using this antibody (Fig. 1f), confirming the specificity of the anti-sfSGK-C antibody.

**Fig. 2.**
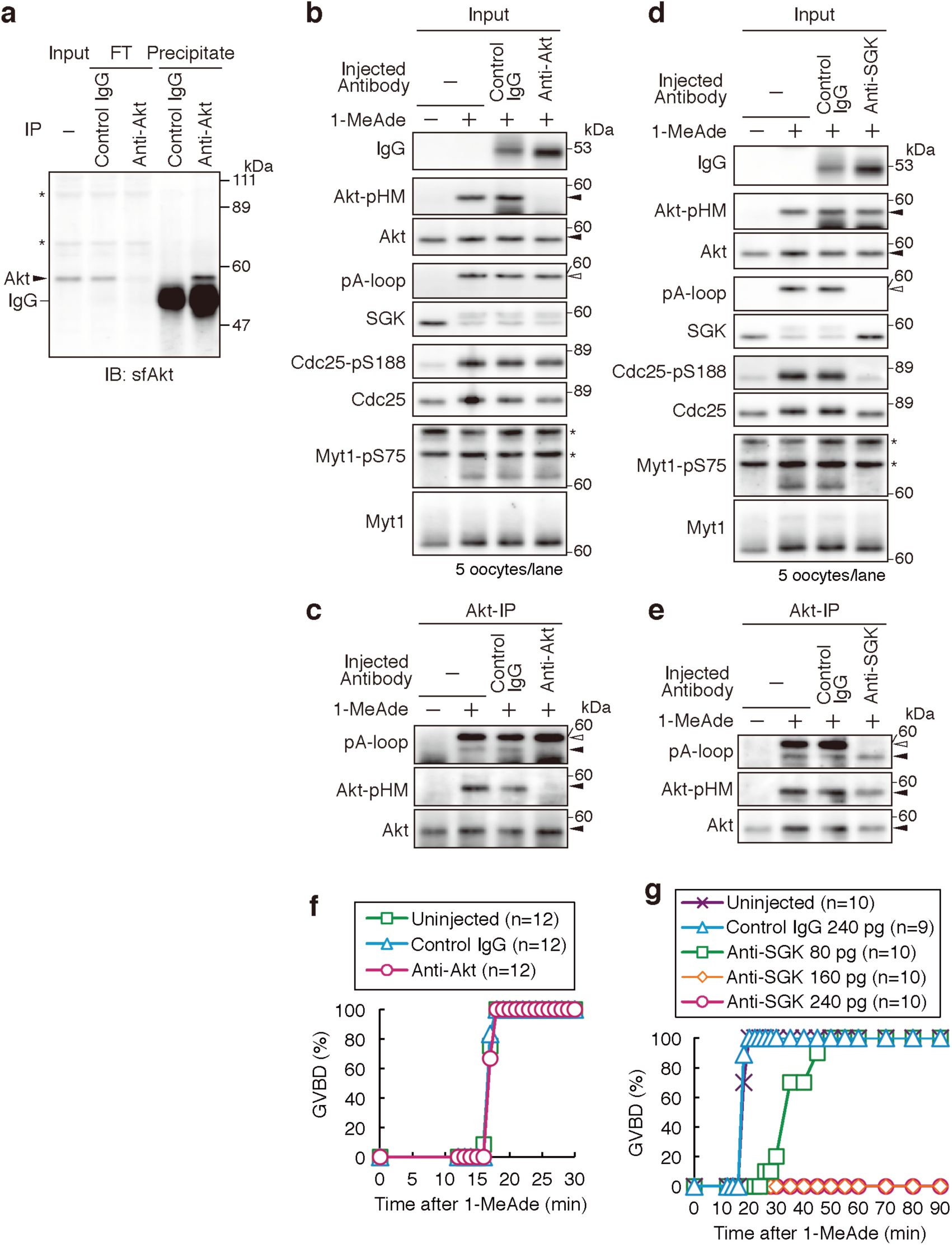
SGK, but not Akt, is required for the meiotic G2/M transition. **a** Immunoprecipitation with control IgG or the anti-sfAkt-C antibody was performed using extracts prepared from unstimulated oocytes, followed by immunoblotting with the anti-sfAkt-C antibody. **b, c** The anti-sfAkt-C antibody inhibits activation of Akt. Unstimulated oocytes were injected with the anti-sfAkt-C antibody or control IgG and then treated with 1-MeAde for 4 min. Immunoprecipitation with the anti-sfAkt-C antibody was performed. Input extracts (**b**, Input) and precipitates (**c**, Akt-IP) were analyzed by immunoblotting. **d, e** The anti-sfSGK-C antibody inhibits activation of SGK. Unstimulated oocytes were injected with the anti-sfSGK-C antibody or control IgG and then treated with 1-MeAde for 4 min. Extracts were prepared, and injected IgG was removed by mixing samples with protein A-Sepharose beads. Thereafter, extracts were subjected to immunoprecipitation with the anti-sfAkt-C antibody. Input extracts (**d**, Input) and precipitates (**e**, Akt-IP) were analyzed by immunoblotting. **f** Akt is not required for the meiotic G2/M transition. Unstimulated oocytes were injected with the anti-sfAkt-C antibody or control IgG and then treated with 1-MeAde. GVBD was monitored as a marker of the meiotic G2/M transition. The graph shows the percentage of oocytes that had undergone GVBD by the indicated time points. “n” indicates the number of oocytes observed. **g** SGK is required for the meiotic G2/M transition. Unstimulated oocytes were injected with the indicated amount of the anti-sfSGK-C antibody or control IgG and then treated with 1-MeAde. GVBD was monitored and plotted as described in **f**. In all panels, open arrowheads indicate the position of A-loop-phosphorylated SGK, closed arrowheads indicate the position of Akt, and asterisks indicate non-specific bands. The data are representative of two independent experiments in **a**-**c**, **f**, and **g**; and three independent experiments in **d** and **e**.

The antigens of these antibodies include the phosphorylation site in the HM of each protein. Thus, the antibodies were expected to bind to the HM and block its phosphorylation. As expected, microinjection of the anti-sfAkt-C antibody abolished HM phosphorylation of Akt after 1-MeAde-stimulation (Fig. 2b). We previously suggested that A-loop phosphorylation of Akt depends on its HM phosphorylation in starfish oocytes based on analysis using exogenously expressed human Akt^47^. Consistently, A-loop phosphorylation was also disrupted by injection of the anti-sfAkt-C antibody (Fig. 2c, closed arrowhead). SGK was activated as normal in these oocytes (Fig. 2b, open arrowhead). Thus, the anti-sfAkt-C antibody specifically inhibits Akt. On the other hand, microinjection of the anti-sfSGK-C antibody disrupted both A-loop phosphorylation and the mobility shift of SGK after 1-MeAde stimulation (Fig. 2d) without affecting phosphorylation of Akt (Fig. 2d for HM, e for A-loop, closed arrowhead), indicating that this antibody specifically inhibits SGK.

We next examined the effect of microinjection of these antibodies on the meiotic G2/M transition. Oocytes were injected with the anti-sfAkt-C antibody and then treated with 1-MeAde. As a marker of the meiotic G2/M transition, we monitored germinal vesicle breakdown (GVBD), which is equivalent to nuclear envelope breakdown in somatic cells. Surprisingly, GVBD occurred as normal in these Akt-inhibited oocytes (Fig. 2f). By contrast, microinjection of the anti-sfSGK-C antibody suppressed GVBD in a dose-dependent manner (Fig. 2g), suggesting that SGK, but not Akt, is required for the meiotic G2/M transition.

We previously reported that microinjection of an anti-sfAkt antibody inhibits the meiotic G2/M transition^39^. The antibody used in that previous study was raised against an 88-amino acid C-terminal fragment of sfAkt. In the present study, there were no stocks of the antiserum left and therefore we used an antibody purified from an antiserum raised against the same fragment of sfAkt in another rabbit. Thus, the specificity of the previously used antibody may differ from that of the presently used antibody. In addition, 33 of the 88 residues in the antigen peptide are conserved in sfSGK and therefore the previously used antibody may have cross-reacted with sfSGK and consequently inhibited SGK as well as Akt. Check of the specificity of this antibody was limited during our previous study because we lacked an anti-sfSGK antibody and phospho-specific antibodies against both sfAkt and sfSGK at that time.

### SGK directly phosphorylates Cdc25 and Myt1 to trigger cyclin B-Cdk1 activation

To trigger activation of cyclin B-Cdk1, we previously demonstrated that Cdc25 and Myt1 are activated and inactivated, respectively, by phosphorylation of Akt/SGK consensus motifs, which correspond to multiple sites including Ser188 in Cdc25 (ref. 48), and Ser75 in Myt1 (ref. 39). Consistent with these previous findings and with the requirement of SGK for the meiotic G2/M transition, phosphorylation of Cdc25 at Ser188 and Myt1 at Ser75 was inhibited by injection of the anti-sfSGK-C antibody (Fig. 2d), but not by injection of the anti-sfAkt-C antibody (Fig. 2b). Furthermore, in Phos-tag SDS-polyacrylamide gel electrophoresis (PAGE) analysis, which emphasizes the mobility shifts of phosphorylated proteins, the upward shifts of Cdc25 and Myt1 after 1-MeAde stimulation were completely inhibited by the anti-sfSGK-C antibody, but not by the anti-sfAkt-C antibody (Fig. 3a). This suggests that the regulatory phosphorylation of Cdc25 and Myt1 to trigger activation of cyclin B-Cdk1 depends on SGK, but not on Akt.

**Fig. 3.**
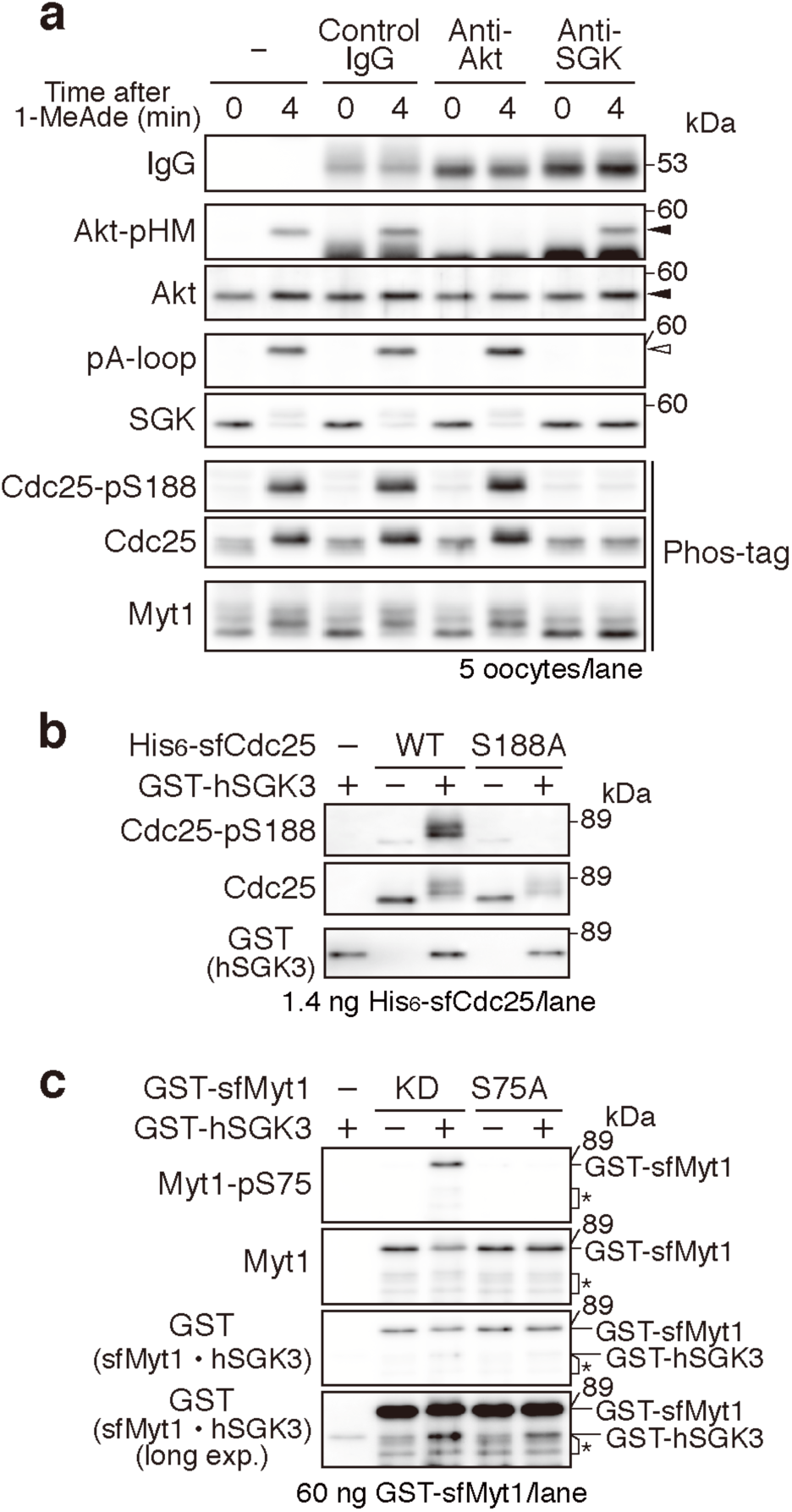
SGK directly phosphorylates Cdc25 and Myt1. **a** SGK, but not Akt, is required for Cdk-independent phosphorylation of Cdc25 and Myt1 upon 1-MeAde stimulation. Unstimulated oocytes were injected with the indicated antibodies, treated with 1-MeAde in the presence of roscovitine, and then analyzed by immunoblotting using a normal or Phos-tag-containing SDS-PAGE gel. The open arrowhead indicates the position of A-loop-phosphorylated SGK, while the closed arrowheads indicate the position of Akt. **b** His-tagged recombinant wild-type (WT) sfCdc25 or the S188A mutant was incubated with or without GST-tagged active human SGK3 (hSGK3) in the presence of ATP and MgCl_2_. Thereafter, immunoblotting was performed. **c** GST-tagged recombinant kinase-dead (KD) or the S75A mutant of sfMyt1 was phosphorylated by human SGK3 as described in **b**. Asterisks indicate minor fragments of recombinant Myt1. The data in all panels are representatives of each two independent experiments.

To investigate whether SGK directly phosphorylates Cdc25 and Myt1, *in vitro* phosphorylation experiments were performed using recombinant proteins. Attempts to prepare active recombinant sfSGK were unsuccessful; therefore, we used commercially available recombinant human SGK3 (hSGK3), which is the best related to sfSGK among the three isoforms of human SGK. Ser188 of starfish Cdc25 (sfCdc25) was phosphorylated upon incubation with hSGK3 (Fig. 3b). In addition, the mobility shift was still observed when Ser188 of sfCdc25 was substituted with Ala (Fig. 3b), suggesting that hSGK3 directly phosphorylates sfCdc25 at multiple residues including Ser188. This is consistent with the fact that sfCdc25 has five Akt/SGK consensus motifs. hSGK3 also phosphorylated Ser75 of starfish Myt1 (sfMyt1), leading to a slight mobility shift (Fig. 3c). Consistent with the fact that sfMyt1 has only one Akt/SGK consensus motif, the mobility shift of sfMyt1 was not observed when Ser75 was substituted with Ala (Fig. 3c), suggesting that Ser75 is the only site phosphorylated by hSGK3 in sfMyt1. These results suggest that Cdc25 and Myt1 are direct substrates for SGK in starfish oocytes.

The importance of SGK in 1-MeAde signaling was further supported by experiments using oocyte extracts. A GST-conjugated Akt/SGK substrate peptide (GST-AS peptide), which contained an Akt/SGK consensus motif, was incubated in oocyte extracts. Thereafter, phosphorylation of the peptide was evaluated by immunoblotting with an anti-pan-phospho-Akt/SGK substrate antibody. The GST-AS peptide was not phosphorylated in the extract of unstimulated oocytes, but was phosphorylated in the extract of 1-MeAde-treated oocytes (Fig. 4a, b). This phosphorylation of the GST-AS peptide was not affected by immunodepletion of Akt (Fig. 4c for confirmation of Akt depletion and d, e for activity), but was abolished upon depletion of SGK (Fig. 4f for confirmation of SGK depletion and g, h for activity). These results suggest that SGK is a major kinase activated after 1-MeAde stimulation and phosphorylates the Akt/SGK consensus motif in starfish oocytes.

**Fig. 4.**
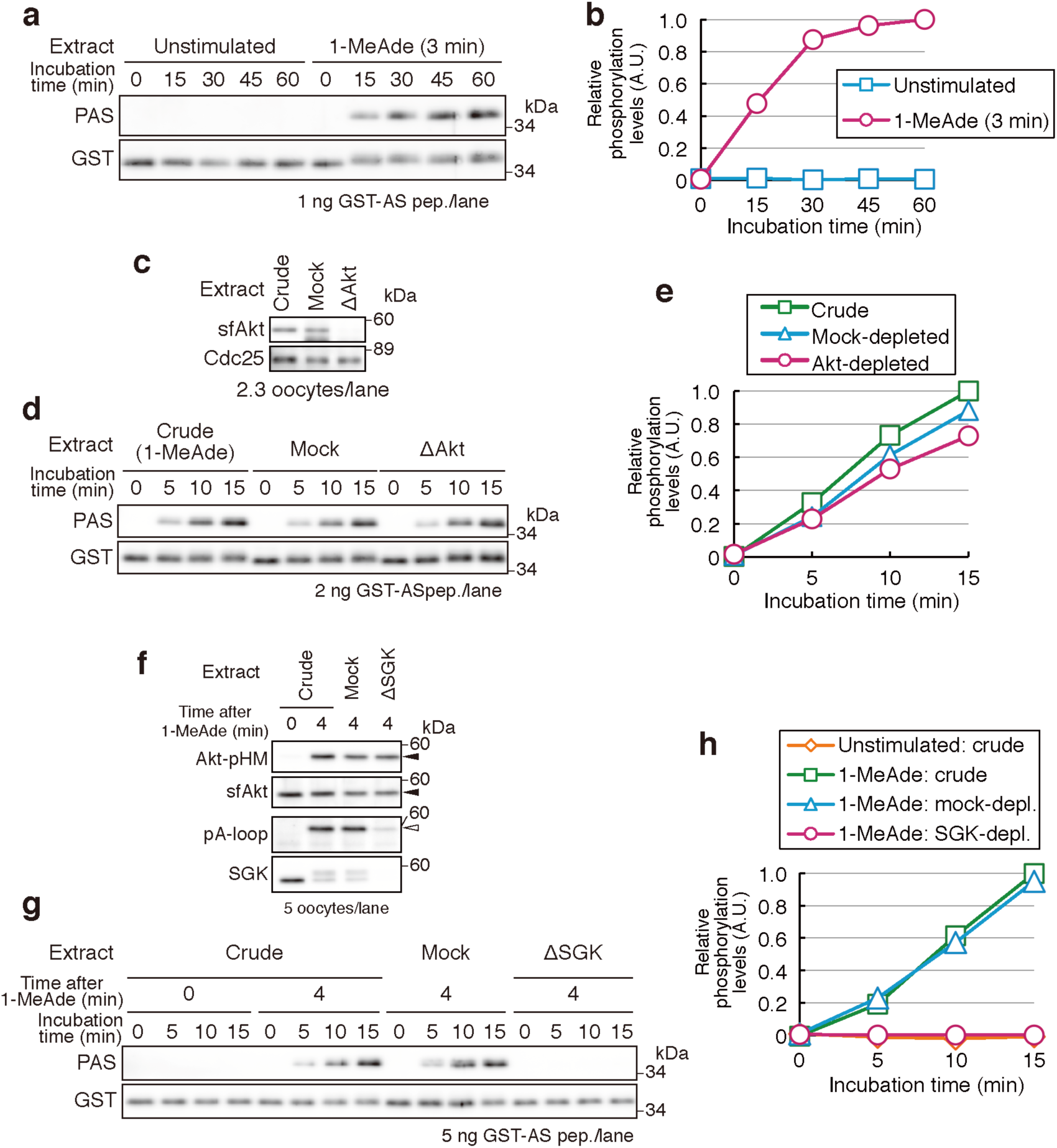
SGK is a major kinase that phosphorylates the Akt/SGK consensus motif in starfish oocytes. **a, b** Kinase activity for the AS peptide is detectable in an extract of 1-MeAde-treated oocytes. Extracts were prepared from unstimulated or 1-MeAde-treated (3 min) oocytes and incubated with the GST-AS peptide for the indicated durations. Phosphorylation of the GST-AS peptide was detected by immunoblotting with an anti-pan-phospho-Akt/SGK substrate antibody (PAS), which recognizes phosphorylated Ser or Thr in the Akt/SGK consensus motif (**a**). Phosphorylation of the GST-AS peptide was quantitated from the immunoblots in **a** and normalized against the total amount of the GST-AS peptide. The graph shows the levels of phosphorylation relative to that at 60 min (**b**). **c–e** Akt is not responsible for the AS-peptide kinase activity in the extract. An extract was prepared from 1-MeAde-treated oocytes. Immunodepletion was performed with control IgG (mock) or the anti-sfAkt-C antibody (ΔAkt). The crude and immunodepleted extracts were analyzed by immunoblotting (**c**). The GST-AS peptide kinase assay was performed using the extracts shown in **c** (**d**). Phosphorylation was quantitated as described in **a** and **b**. The graph shows the levels of phosphorylation relative to that at 15 min in the crude extract (**e**). **f–h** SGK is responsible for the AS-peptide kinase activity in the extract. An extract was prepared from unstimulated or 1-MeAde-treated oocytes. Immunodepletion was performed with the anti-sfSGK-C antibody (**f**), the GST-AS peptide kinase assay was performed (**g**), and phosphorylation was quantitated (**h**) as described in **c**, **d**, and **e**. The open arrowhead indicates the position of A-loop-phosphorylated SGK, while the closed arrowheads indicate the position of Akt. The data in all panels are representatives of each two independent experiments.

Taken together, these observations support the conclusion that SGK, but not Akt, is required for the regulatory phosphorylation that makes Cdc25 activity dominant over Myt1 activity to trigger cyclin B-Cdk1 activation.

### SGK is sufficient for the regulatory phosphorylation of Cdc25 and Myt1 to trigger cyclin B-Cdk1 activation at the meiotic G2/M transition

Next, we investigated whether sfSGK is sufficient to trigger cyclin B-Cdk1 activation. Mammalian SGK3 and Akt have a regulatory domain N-terminal to their catalytic domain, which is named the PX domain and PH domain, respectively^51^. Binding of these domains to phosphoinositides on the cellular membrane contributes to activation of these kinases^55,56^. Membrane targeting of Akt via substitution of the PH domain with a myristoylation sequence results in its activation without extracellular stimulation^39,47,57^. To generate constitutively active sfSGK (CA-SGK), we substituted the PX domain of sfSGK with a myristoylation sequence. Furthermore, Thr479, which is the conserved phosphorylation site by TORC2, was substituted with Glu to mimic phosphorylation of the HM. Injection of mRNA encoding CA-SGK into unstimulated oocytes induced GVBD (Fig. 5a). In these oocytes, the level of A-loop phosphorylation of CA-SGK at GVBD was comparable to that of endogenous SGK after 1-MeAde stimulation (Fig. 5b, c), suggesting that CA-SGK is activated in the absence of 1-MeAde stimulation and that activation of SGK is sufficient to induce the meiotic G2/M transition.

**Fig. 5.**
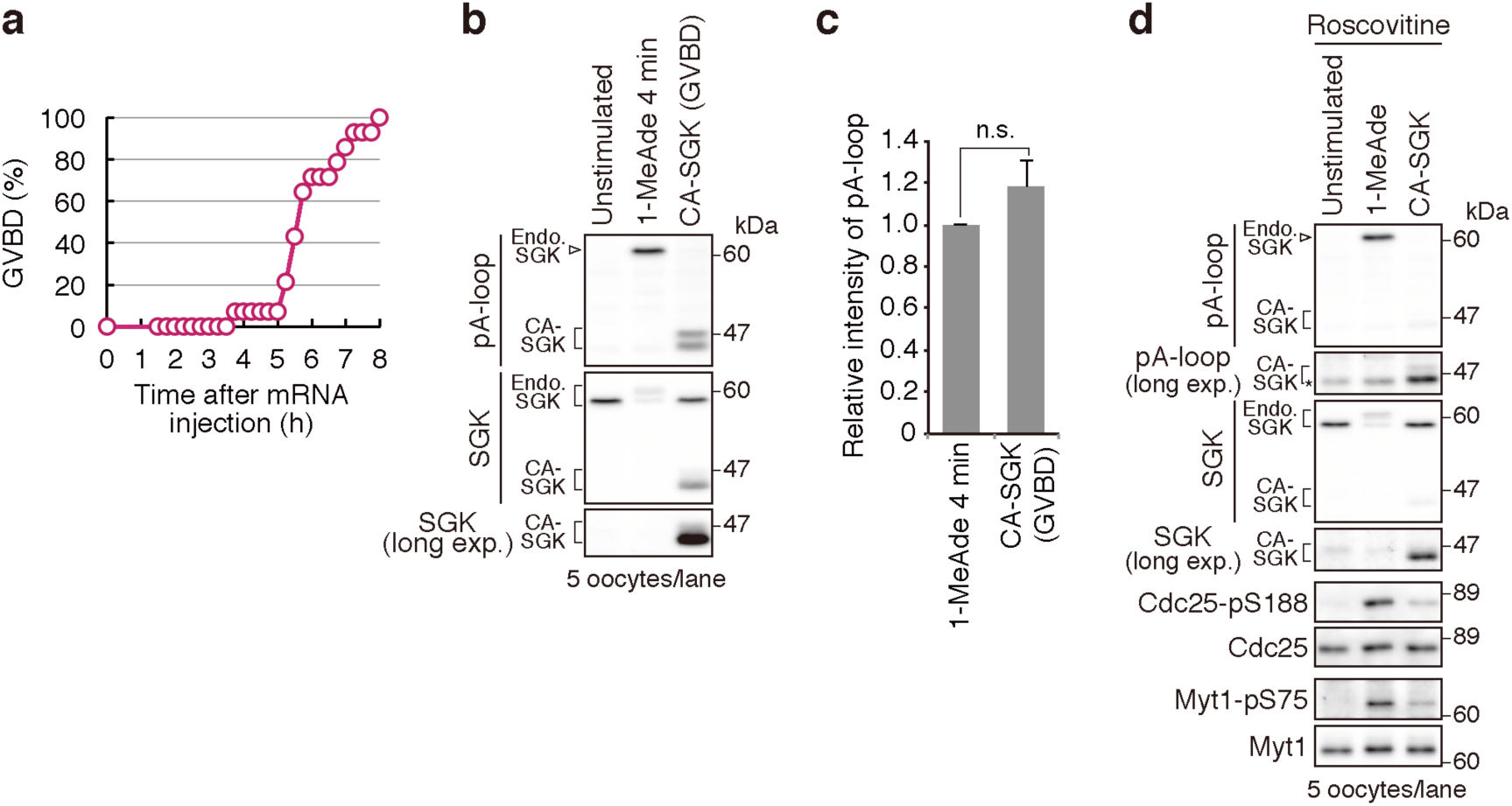
SGK is sufficient for the meiotic G2/M transition. **a–c** CA-SGK expression induces the meiotic G2/M transition. Unstimulated oocytes were injected with mRNA encoding CA-SGK and incubated. GVBD in 14 oocytes was monitored every 15 min. The graph shows the percentage of oocytes that had undergone GVBD by the indicated time points (**a**). CA-SGK-expressing oocytes at GVBD, unstimulated oocytes, and 1-MeAde-treated oocytes were analyzed by immunoblotting (**b**). Phosphorylation levels of the A-loop of endogenous SGK in 1-MeAde-treated oocytes and CA-SGK at GVBD were quantitated from the immunoblots in **b**. The graph shows the relative phosphorylation levels (**c**). Data represent mean values ± standard deviation from three independent experiments. P = 0.06 (one-tailed t-test; not significant, n.s.). **d** CA-SGK expression induces regulatory phosphorylation of Cdc25 and Myt1 to trigger cyclin B-Cdk1 activation. Unstimulated oocytes were treated with 1-MeAde for 4 min in the presence of roscovitine. For expression of CA-SGK, unstimulated oocytes were injected with the mRNA, incubated with roscovitine, collected at the time point at which 50% of CA-SGK-expressing oocytes exhibited GVBD in the absence of roscovitine (5 h 45 min in **a**), and analyzed by immunoblotting. Note that, in the immunoblot of phospho-A-loop after a long exposure (pA-loop, long exp.), weak non-specific bands were detected at almost the same size as CA-SGK (asterisk). The data are representative of three independent experiments in **a**, **b**, and **c**; and two independent experiments in **d**.

We then examined whether CA-SGK expression induces the regulatory phosphorylation of Cdc25 and Myt1 to trigger cyclin B-Cdk1 activation. To analyze CA-SGK-induced signaling without any effect of Cdk-dependent feedback pathways, CA-SGK was expressed in the presence of the Cdk inhibitor roscovitine. These oocytes were collected at the time point at which 50% of CA-SGK-expressing oocytes exhibited GVBD in the absence of roscovitine (5 h and 45 min in Fig. 5a). Although expression and A-loop phosphorylation levels of CA-SGK were lower than those of endogenous SGK (Fig. 5d, pA-loop), phosphorylation of Cdc25 at Ser188 and of Myt1 at Ser75 were detectable in these oocytes (Fig. 5d). Taken together, these findings suggest that activation of SGK is sufficient for the regulatory phosphorylation of Cdc25 and Myt1 to trigger cyclin B-Cdk1 activation.

### SGK is activated by cooperation of the Gβγ-PI3K and atypical Gβγ pathways in starfish oocytes

In a previous report^48^, we proposed the existence of an atypical Gβγ pathway that is activated by Gβγ and cooperates with the Gβγ-PI3K pathway to induce the meiotic G2/M transition. Therefore, we examined whether SGK is activated by these pathways.

First, we investigated whether SGK is activated by signaling pathways downstream of Gβγ. Expression of exogenous Gβγ induced the meiotic G2/M transition as previously reported^48^, whereas Gβγ-induced GVBD was inhibited by microinjection of the anti-sfSGK-C antibody, but not by injection of the anti-sfAkt-C antibody (Fig. 6a). These findings indicate that signaling from exogenous Gβγ induces the meiotic G2/M transition in a SGK-dependent manner, similar to 1-MeAde stimulation. Consistently, immunoblot analysis showed that SGK was activated upon expression of Gβγ (Fig. 6b). Furthermore, when Gβγ was expressed in the presence of the Cdk inhibitor roscovitine, thereby avoiding any effect of cell cycle progression on signal transduction, the level of A-loop phosphorylation of sfSGK was comparable to that induced by 1-MeAde (Fig. 6c), suggesting that sfSGK is activated downstream of Gβγ independently of Cdk activity. However, 1-MeAde- and Gβγ-induced SGK activation was abolished upon inhibition of PI3K by wortmannin (Fig. 6d), suggesting that activation of sfSGK by Gβγ requires PI3K.

**Fig. 6.**
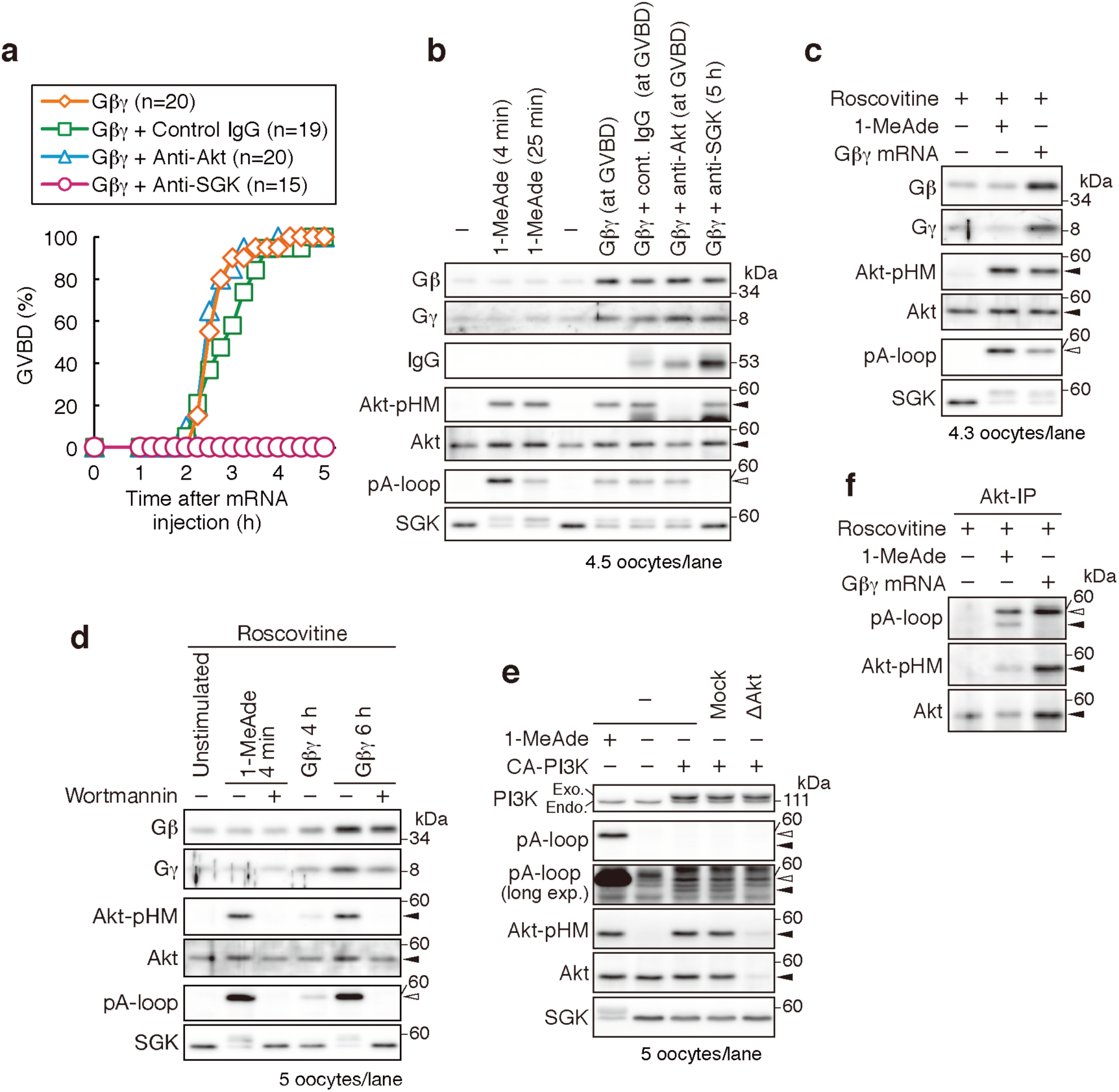
SGK is activated by cooperation of the Gβγ-PI3K and atypical Gβγ pathways. **a, b** Exogenous expression of Gβγ induces the meiotic G2/M transition in a SGK-dependent manner. Unstimulated oocytes were injected with the indicated antibodies and subsequently injected with mRNA encoding Gβγ. Thereafter, GVBD was monitored (**a**). Oocytes were collected when GVBD occurred. Anti-sfSGK antibody-injected oocytes, which did not undergo GVBD, were collected at 5 h after mRNA injection. For 1-MeAde treatment, unstimulated oocytes were treated with 1-MeAde for 4 or 25 min (GVBD occurred at approximately 21 min) and then analyzed by immunoblotting (**b**). **c** Exogenous Gβγ expression induces SGK activation. Unstimulated oocytes were treated with 1-MeAde for 4 min in the presence of roscovitine or injected with mRNA encoding Gβγ and then incubated with roscovitine for 6 h. Thereafter, extracts were prepared from these oocytes and analyzed by immunoblotting. These extracts were also used for immunoprecipitation of Akt in **f**. **d** Gβγ-induced SGK activation requires PI3K. Unstimulated oocytes were treated with 1-MeAde or injected with mRNA encoding Gβγ, incubated in the presence of roscovitine with or without wortmannin, and analyzed by immunoblotting. **e** PI3K is not sufficient for activation of SGK. Unstimulated oocytes were treated with 1-MeAde for 4 min or injected with mRNA encoding CA-PI3K and incubated for 3 h. Extracts of CA-PI3K-expressing oocytes were immunodepleted using control IgG (Mock) or the anti-sfAkt-C antibody (ΔAkt). Thereafter, immunoblotting was performed. **f** Expression of exogenous Gβγ fails to induce A-loop phosphorylation of Akt. Endogenous Akt was immunoprecipitated from the extracts in **c** and analyzed by immunoblotting. In all panels, open arrowheads indicate the position of A-loop-phosphorylated SGK, while closed arrowheads indicate the position of Akt. The data are representative of two independent experiments in **a**, **b**, **d**, and **e**; and three independent experiments in **c** and **f**.

Next, to determine whether PI3K activation solely induces SGK activation, CA-PI3K was expressed in unstimulated oocytes. As previously reported^48^, CA-PI3K induced HM phosphorylation of Akt (Fig. 6e), but not the meiotic G2/M transition. In these oocytes, A-loop phosphorylation of SGK was only detectable by immunoblotting with a long time exposure for acquisition of the chemiluminescent image (Fig. 6e, pA-loop with long exposure, open arrowhead), indicating that activation of PI3K barely induces activation of SGK.

Taken together, Gβγ activates SGK in a PI3K-dependent manner, but activation of PI3K alone is insufficient for SGK activation. These observations suggest that Gβγ activates not only PI3K but also another pathway(s), which we call the atypical Gβγ pathway, and that cooperation of the Gβγ-PI3K and atypical Gβγ pathways leads to activation of SGK.

As to activation of Akt, CA-PI3K induced HM phosphorylation of Akt (Fig. 6e), as previously reported^48^. In these oocytes, the anti-phospho-A-loop antibody detected a weak band at the same mobility as Akt by immunoblotting after a long exposure (Fig. 6e, pA-loop with long exposure, closed arrowhead) even in whole-oocyte samples. This band disappeared after immunodepletion of Akt from the oocyte extract (Fig. 6e), showing that it corresponds to A-loop phosphorylation of Akt. Given that the level of 1-MeAde-induced A-loop phosphorylation of Akt was below the threshold of detection in whole-oocyte samples (Fig. 1b–d), Akt is activated more strongly by CA-PI3K than by 1-MeAde. This further indicates that activation of endogenous Akt is insufficient for the meiotic G2/M transition. By contrast, in Gβγ-expressing oocytes, A-loop phosphorylation of Akt was undetectable even in immunoprecipitated Akt samples (Fig. 6f, closed arrowhead), although HM phosphorylation was comparable to that in 1-MeAde-treated oocytes (Fig. 6c, Akt-pHM in Input). This suggests that Gβγ signaling is insufficient for A-loop phosphorylation of Akt. This finding will be discussed later.

## Discussion

This study showed that SGK is necessary and sufficient for the regulatory phosphorylation of Cdc25 and Myt1 to trigger activation of cyclin B-Cdk1 in the meiotic G2/M transition. Furthermore, we revealed that SGK is activated by cooperation of the Gβγ-PI3K and atypical Gβγ pathways downstream of 1-MeAde stimulation (Fig. 7). These findings clarify the molecular identity of the trigger kinase, and offer important clues to elucidate the 1-MeAde signaling pathway. Moreover, they provide new insights into the role and regulation of SGK.

**Fig. 7.**
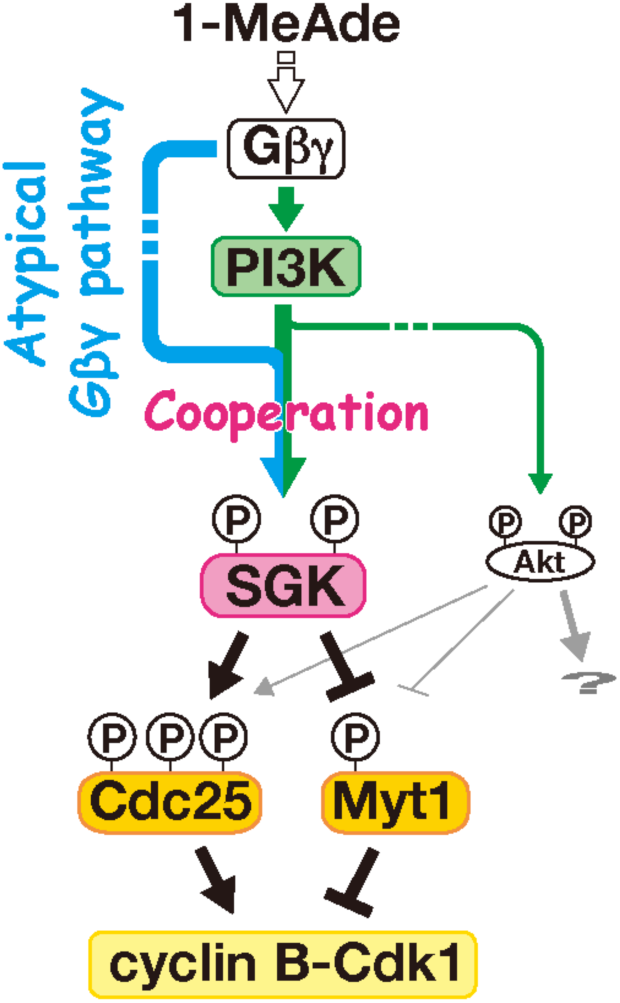
SGK triggers cyclin B-Cdk1 activation in 1-MeAde signaling. Downstream of 1-MeAde stimulation, Gβγ activates the PI3K pathway (green arrow) and the atypical Gβγ pathway (cyan arrow). SGK is activated by cooperation of these pathways and subsequently activates and inactivates Cdc25 and Myt1, respectively, via direct phosphorylation. Consequently, reversal of the balance of Cdc25 and Myt1 activities triggers cyclin B-Cdk1 activation. Akt is weakly activated after 1-MeAde stimulation, and may phosphorylate Cdc25 and Myt1 at undetectable levels. However, Akt is dispensable for cyclin B-Cdk1 activation and may have any other function in oocytes.

In *Xenopus* oocytes, multiple pathways are proposed to reverse the balance between the activities of Cdc25 and Myt1 (refs 23,30). Therefore, it is difficult to determine the relative contribution of each pathway. In this context, SGK is the only trigger kinase in starfish oocytes because it is necessary and sufficient for the regulatory phosphorylation of Cdc25 and Myt1. Particularly, the phosphorylation of Cdc25 and Myt1 was undetectable in SGK-inhibited oocytes (Fig. 2). In mouse oocytes, Cyclin B-Cdk1 activation might be triggered by a phosphatase antagonistic to PKA, rather than by kinases, and it is unclear whether the activity of this phosphatase is regulated at the meiotic G2/M transition^29^. Thus, our present study is the first time that the trigger kinase in oocyte meiosis has been clearly identified. To the best of our knowledge, this is the first report that SGK is involved in M-phase entry.

In parallel with the present study, Hosoda et al. showed that sfSGK is required for the increase in intracellular pH after 1-MeAde stimulation in starfish oocytes^54^. They suggested that the increase in intracellular pH is not involved in cyclin B-Cdk1 activation, but is required for progression of GVBD after full activation of cyclin B-Cdk1 in ovarian oocytes. The functions of SGK have been mainly studied in mammalian cells, which express three SGK isoforms: SGK1, SGK2, and SGK3^51^. Although mammalian SGKs are reportedly involved in various processes, such as cell survival, stress responses, and tumorigenesis, in somatic cells^58,59^, their functions in germ cells remain unclear. To the best of our knowledge, studies presented here and in the paper by Hosoda et al.^54^ are the first indication of the functions of SGK in germ cells.

Among the three mammalian SGK isoforms, only SGK3 contains a regulatory domain, called the PX domain, N-terminal to the kinase domain^51,60^. sfSGK also has a PX domain and is therefore likely an ortholog of mammalian SGK3 (ref. 54). The PX domain of mammalian SGK3 binds to phosphatidylinositol 3-phosphate (PI-3P), which is a phosphorylated derivative of phosphatidylinositol (PI) at the 3-position of the inositol ring^56^. This interaction is essential for the localization and activation of SGK3 (refs 50,56). PI-3P is generated by dephosphorylation of PI-345P_3_ at the 4- and 5-positions of the inositol ring^50^. PI3K, which produces PI-345P_3_ from PI-45P_2_, is activated in signaling pathways downstream of receptor tyrosine kinases and GPCRs^45^. Thus, production of PI-3P from PI-345P_3_ may occur downstream of these receptors. Indeed, a recent report suggested that this mechanism is implicated in activation of SGK3 after stimulation by IGF-1, which is an agonist for a receptor tyrosine kinase^61^. In this context, SHIP2 and INPP4A/B were proposed to be phosphatases for PI-3P production, although the regulatory mechanisms remain elusive^61^. Furthermore, it is unclear whether PI-345P_3_ is a source of PI-3P for SGK activation in GPCR-dependent signaling. We previously reported that PI-345P_3_ is generated on the plasma membrane after 1-MeAde stimulation in starfish oocytes^48^. This PI-345P_3_ may be metabolized to PI-3P for SGK activation. Our present results suggest that the atypical Gβγ pathway cooperates with PI3K to activate sfSGK (Fig. 6). Thus, we propose that the atypical Gβγ pathway activates phosphatases that produce PI-3P from PI-345P_3_ downstream of GPCR-mediated signaling.

Another mechanism for PI-3P production is phosphorylation of PI at the 3-position, which occurs on early endosomes during intracellular vesicle trafficking and endocytosis^45,50^. GPCRs are frequently endocytosed after agonist stimulation^62^. Thus, 1-MeAde may induce endocytosis, which increases the level of early endosomes. If this is the case, the endocytic pathway may also contribute to activation of SGK in starfish oocytes. Many aspects of these scenarios, such as the dynamics of PI-3P, the localization of SGK, and the molecular identity of the atypical Gβγ pathway, remain unclear. Nonetheless, these working models could be tested in future studies and may help to elucidate the regulatory mechanisms of SGK3 activation in mammalian somatic cells.

SGKs and Akt reportedly play redundant roles in some contexts^59^. In starfish oocytes, Akt has been reported to have a potential to trigger cyclin B-Cdk1 activation^39^. Indeed, an excess amount of CA-Akt expression can induce the meiotic G2/M transition^39,48^. Furthermore, regulatory effects of phosphorylation of the Akt/SGK consensus motifs on Myt1 and Cdc25 activities were originally showed using Akt as a kinase in *in vitro* experiments^39,48^. Nonetheless, our present results showed that SGK is predominantly responsible for these phosphorylation *in ovo*. Contribution of Akt was, if any, undetectable (Fig. 2) possibly due to its weak activation, as supported by low levels of A-loop phosphorylation (Figs. 1) and by the kinase assay using oocyte extracts (Fig. 5). The role of Akt may be other than the triggering of cyclin B-Cdk1 activation. Even so, we cannot exclude the possibility that Akt makes a significant contribution to the triggering as a redundant kinase with SGK in a limited number of unknown circumstances in which Akt is more abundant than SGK in oocytes.

Contrary to our present conclusion, we previously reported that Akt is necessary for the meiotic G2/M transition^39^. In our previous report, in addition to the anti-sfAkt antibody, we also used competitive peptides derived from the A-loop or HM of Akt to inhibit Akt activation. Akt and SGK have common upstream kinases; therefore, peptide competition likely inhibited activation of both Akt and SGK. In the present study, successful generation of anti-phospho-A-loop and anti-sfSGK-C antibodies enabled us to evaluate the specificities of the neutralizing antibodies more precisely. In addition, Akt was dispensable at least in all cases tested in the current study, which used oocytes isolated from six starfish collected at three different areas in Japan. Thus, our current conclusions are more convincing than our previous conclusions.

We made some interesting observations regarding activation of Akt. Distinct from SGK, activation of Akt depends on binding of PI-345P_3_ to its PH domain^59^. Expression of CA-PI3K, but not of Gβγ, induced A-loop phosphorylation of Akt (Fig. 6e, f). We previously reported that a comparable level of PI-345P_3_ is produced upon 1-MeAde stimulation and Gβγ expression, while CA-PI3K induces a higher level of PI-345P_3_ (ref. 48). Thus, it is possible that a high level of PI-345P_3_ alone can induce Akt activation, whereas a low level of PI-345P_3_ requires an additional mechanism to enhance A-loop phosphorylation. In addition, it should be noted that the levels of HM phosphorylation induced by CA-PI3K and exogenous Gβγ were comparable to that induced by 1-MeAde stimulation (Fig. 6c, e). Thus, the threshold level of PI-345P_3_ required for HM phosphorylation may be lower than that required for A-loop phosphorylation.

In summary, we identified SGK as the trigger kinase for activation of cyclin B-Cdk1 at M-phase entry and as the target of the collaborative actions of PI3K and the atypical Gβγ pathway. These findings not only demonstrate the novel role of SGK in the triggering of M-phase, but also provide insights into the regulatory mechanisms of SGK activation, particularly with regard to PI-3P production.

## Methods

### Chemicals

Roscovitine (Calbiochem), wortmannin (LC Laboratories), and BX795 (Enzo Life Sciences) were dissolved in DMSO at concentrations of 50, 20, and 6 mM, respectively, as stock solutions and used at final concentrations of 45, 40, and 6 µM, respectively.

### DNA constructs

For CA-SGK, an N-terminal fragment of sfSGK containing the PX domain (Met1–Asp132) was substituted with a myristoylation sequence (MGSSKSKPKDPSQR) by PCR. Thereafter, the insert was cloned into a modified pSP64-S vector, in which the SP6 promoter for *in vitro* transcription had been substituted by a T7 promoter sequence^48,63^, using an In-Fusion kit (Takara Bio). Finally, Thr479 in the HM was substituted with Glu by PCR, and the point mutation was verified by sequencing. Constructs encoding untagged sfAkt, CA-PI3K in which Flag tag and plasma membrane-targeting sequence including a CAAX motif fused to the N- and C-terminus of the p110 catalytic subunit of sfPI3K, respectively, untagged starfish Gβ (sfGβ), and untagged starfish Gγ (sfGγ) for mRNA preparation were prepared as described previously^48^.

### Preparation of recombinant proteins

Recombinant sfMyt1 (N229A for the kinase-dead mutant or S75A mutant) cloned into the pGEX-4T-1 vector was expressed in *Escherichia coli* (*E. coli*) strain BL21 (DE3) (Invitrogen) as an N-terminal GST-tagged protein, purified using glutathione-Sepharose 4B beads (GE Healthcare), and dialyzed against storage buffer (20 mM PIPES pH 6.8, 200 mM sucrose, and 1 mM DTT). For His_6_-Cdc25 (wild-type or S188A mutant), *E. coli* strain BL21 (DE3) was transformed with the pET21a construct. Expressed His_6_-Cdc25 were purified from inclusion bodies under denaturing conditions (6 M urea) using Probond-Resin (Invitrogen) and then refolded via stepwise reduction of the urea concentration by dialysis using EasySep (TOMY) for 2 h against Tris-buffered saline (TBS; 50 mM Tris and 150 mM NaCl, pH 7.5) containing 4 M urea, for 2 h against TBS containing 2 M urea, and for 2 h against TBS lacking urea. To prepare the recombinant GST-fused AS peptide (AGRPRAATFIESG), *E. coli* strain BL21-CodonPlus-RIL (Agilent Technologies) was transformed with the pGEX-4T-1 construct. The expressed GST-AS peptide was purified using glutathione-Sepharose 4B and dialyzed against phosphate-buffered saline (137 mM NaCl, 2.68 KCl, 10 mM Na_2_HPO_4_, and 1.76 mM KH_2_PO_4_, pH 7.4). A recombinant C-terminal 88-amino acid fragment of sfAkt was prepared as previously described^39^.

### Antibody generation

Antibodies were generated by immunizing rabbits (Biologica) and then purified using antigens. Antigens used for immunization and antibody purification were a phospho-peptide of the A-loop of sfAkt (Thr304–Pro320, phosphorylated at Thr315) for the anti-phospho-A-loop antibody, a C-terminal peptide of sfSGK (Ser473–Asp489) for the anti-sfSGK-C antibody^54^, and a C-terminal 88-amino acid fragment of sfAkt for the anti-sfAkt-C antibody. Control IgG for microinjection or immunoprecipitation was purified from rabbit pre-immune serum using protein A-Sepharose 4B (Sigma).

### Oocyte preparation

Starfish *Asterina pectinifera* (renamed *Patiria pectinifera* in the 2007 National Center for Biotechnology Information Taxonomy Browser) was collected during the breeding season and kept in laboratory aquaria supplied with circulating seawater at 14°C. Fully grown unstimulated oocytes with follicle cells were manually isolated from ovaries using forceps. Thereafter, the follicle cells were removed by washing oocytes with calcium-free artificial seawater (476 mM NaCl, 10 mM KCl, 36 mM MgCl_2_, 18 mM MgSO_4_, and 20 mM H_3_BO_3_, pH 8.2). All treatments with 1-MeAde or other chemicals, as well as microinjections, were performed in artificial seawater (ASW; 462 mM NaCl, 9 mM CaCl_2_, 10 mM KCl, 36 mM MgCl_2_, 18 mM MgSO_4_, and 20 mM H_3_BO_3_, pH 8.2) at 23°C. Oocytes were stimulated with 0.5 µM 1-MeAde.

### Immunoblotting

Oocytes were lysed by vortexing in Laemmli sample buffer (LSB) and then heated at 95°C for 5 min. Proteins were separated by SDS-PAGE^64^ on an 8% or 8.5% gel (separating gel: 8% or 8.5% acrylamide, 0.22% or 0.23% N,N’-methylenebisacrylamide, 375 mM Tris, 0.1% SDS, 0.1% APS, and 0.1% TEMED, pH 8.8; stacking gel: 3.6% acrylamide, 0.096% N,N’-methylenebisacrylamide, 120 mM Tris, 0.1% SDS, 0.2% APS, and 0.1% TEMED, pH 6.8; and electrophoresis buffer: 25 mM Tris, 192 mM glycine, and 0.1% SDS) and transferred to a PVDF membrane (Millipore)^65^ using transfer buffer (100 mM Tris, 192 mM glycine, 0.1% SDS, and 20% methanol). Phos-tag SDS-PAGE was performed in accordance with the manufacturer’s protocol using an 8% polyacrylamide gel containing 4 µM Phos-tag (FUJIFILM) and 8 µM MnCl_2_. A modified 15% gel and buffer were used to detect Gγ (separating gel: 15% acrylamide, 0.41% N,N’-methylenebisacrylamide, 750 mM Tris, 0.1% SDS, 0.1% APS, and 0.1% TEMED, pH 8.8; stacking gel: 3.6% acrylamide, 0.096% N,N’-methylenebisacrylamide, 240 mM Tris, 0.1% SDS, 0.2% APS, and 0.1% TEMED, pH 6.8; and electrophoresis buffer: 50 mM Tris, 384 mM glycine, and 0.1% SDS). The membranes were then blocked by incubation in blocking buffer (5% skimmed milk prepared in TBS containing 0.1% Tween-20 (TBS-T)) for 1 h for the anti-phospho-sfMyt1-Ser75, anti-sfMyt1, and anti-GST antibodies, but not for the other antibodies. The primary antibodies were anti-phospho-A-loop (purified, 1:50 in TBS-T), anti-sfAkt-C (purified, 1:1,000 in Can Get Signal Solution 1), anti-phospho-sfAkt-Ser477 (ref. 47, purified, 1:1,000 in Can Get Signal Solution 1), anti-sfSGK-C (purified, 1:2,000 in Can Get Signal Solution 1), anti-phospho-sfCdc25-Ser188 (ref. 48, serum, 1:1,000 in Can Get Signal Solution 1), anti-sfCdc25 (ref. 33, serum, 1:2,000 in Can Get Signal Solution 1), anti-phospho-Cdk1-Tyr15 (Cell Signaling Technology, 1:1,000 in Can Get Signal Solution 1), anti-PSTAIR^47^ to detect Cdk1 (1:50,000 in Can Get Signal Solution 1), anti-phospho-sfMyt1-Ser75 (ref. 48, purified, 1:100 in blocking buffer), anti-sfMyt1^39^ (purified, 1:200 in TBS-T), anti-GST (GE Healthcare, 1:500 in blocking buffer), anti-pan-phospho-Akt/SGK substrate (Cell Signaling Technology, 1:1,000 in Can Get Signal Solution 1), anti-sfp110β to detect sfPI3K^48^ (purified, 1:1,000 in Can Get Signal Solution 1), anti-sfGβ^48^ (serum, 1:1,000 in Can Get Signal Solution 1), and anti-sfGγ^48^ (purified, 1:100 in Can Get Signal Solution 1). Horseradish peroxidase-conjugated secondary antibodies were anti-goat IgG (Sigma, 1:500 in blocking buffer for anti-GST antibody), anti-mouse IgG (Dako, 1:2,000 in Can Get Signal Solution 2 for anti-PSTAIR antibody), and anti-rabbit IgG (GE Healthcare; 1:2,000 in TBS-T for anti-phospho-A-loop, anti-sfSGK-C, anti-phospho-sfMyt1-Ser75, and anti-pan-phospho-Akt/SGK substrate antibodies; in blocking buffer for the anti-sfMyt1 antibody; and in Can Get Signal Solution 2 for the other antibodies). When the signal derived from IgG heavy chains hampered detection, TrueBlot (ROCKLAND, 1:2,000 in Can Get Signal Solution 2) was used instead of the secondary antibody. Signals were visualized with ECL Prime (GE Healthcare), and digital images were acquired on a LAS4000 mini imager (Fujifilm). Quantitation was performed using ImageJ (National Institutes of Health, USA). Graphs were generated using Microsoft Excel. Brightness and contrast were adjusted using ImageJ.

### Microinjection

Microinjection was performed as described previously^66^. mRNAs for exogenous protein expression in starfish oocytes were transcribed from pSP64-S constructs using a mMESSAGE mMACHINE kit (Ambion), dissolved in water, and injected into unstimulated oocytes (20 pg for sfAkt, 20 pg for CA-SGK, and 40 pg for CA-PI3K). To express starfish Gβγ, 50 pg of an equimolar mixture of mRNAs encoding sfGβ and sfGγ was injected. The incubation time for protein expression was determined based on the amount of time taken to induce GVBD or to induce HM phosphorylation of Akt to a comparable extent as that induced by 1-MeAde. For microinjection of antibodies, concentration of antibodies and buffer exchange with phosphate-buffered saline were performed using a 50 k cutoff Amicon Ultra filter (Millipore). Nonidet P-40 (NP-40) was added at a final concentration of 0.05%. Unstimulated oocytes were injected with 230 pg of the anti-TOR antibody, 85 pg of the anti-sfAkt-C antibody, or 200 pg of the anti-sfSGK-C antibody. 1-MeAde was added after incubation for 1 h.

### Immunoprecipitation and immunodepletion

To prepare antibody-bound beads, antibodies (anti-sfAkt antibody, 2.5 µg/µl beads, 0.83 µg for 10 oocytes; and anti-sfSGK antibody, 2 µg/µl beads, 2.7 µg for 10 oocytes) were incubated with Protein A-Sepharose (Sigma) for more than 2 h on ice. Antibody-bound beads were washed with TBS and lysis buffer (80 mM β-glycerophosphate, 20 mM EGTA, 10 mM MOPS pH 7.0, 100 mM sucrose, 100 mM KCl, 1 mM DTT, 1× cOmplete EDTA-free (Merck), 0.5 mM sodium orthovanadate, 1 µM okadaic acid (Calbiochem), and 0.5% NP-40) and used for immunoprecipitation. For cross-linking, antibody-bound beads were washed with TBS and borate buffer (0.2 M NaCl and 0.2 M boric acid pH 9.0) and then incubated with 3.75 mM disuccinimidyl suberate prepared in borate buffer at 24°C for 30 min. Thereafter, the supernatant was discarded and the beads were further incubated in 1 M ethanolamine (pH 8.0) at 24°C for 15 min and finally washed with TBS and lysis buffer. For immunoprecipitation, freeze-thawed oocytes in ASW (10 oocytes/µl of ASW for immunoprecipitation of Akt and SGK) were lysed by adding 6 volumes of lysis buffer and incubating samples on ice for 30 min, followed by gentle vortexing. Lysates were centrifuged at 20 k × g at 4°C for 15 min. The supernatant represented the oocyte extract. Antibody-bound beads were incubated with the oocyte extract (60 oocytes for immunoprecipitation of Akt, 30 oocytes for depletion of Akt, and 15 oocytes for immunoprecipitation and depletion of SGK) for 90 min on ice, separated from flow-through extracts, and washed with lysis buffer. For immunoprecipitation of Akt from oocytes that had been injected with the anti-sfSGK antibody, extracts were pre-incubated with protein A-Sepharose 4B to remove injected IgG. LSB was added to the input extracts, flow-through extracts, and beads for immunoblot analysis. All samples were then heated at 95°C for 5 min and subjected to SDS-PAGE. To prepare mock-, Akt-, and SGK-depleted extracts for the GST-AS peptide kinase assay, freeze-thawed oocytes in ASW (5 oocytes/µl of ASW) were lysed by adding 6.5 volumes of lysis buffer. The extract was then prepared as described above and mixed with beads bound to control IgG, the anti-sfAkt-C antibody, or the anti-sfSGK-C antibody. After incubation at 4°C for 45 min, the supernatant was analyzed by immunoblotting or used for the GST-AS peptide kinase assay.

### *In vitro* phosphorylation of sfCdc25 and sfMyt1

Recombinant sfCdc25 (final concentration of 5 ng/µl) or sfMyt1 (final concentration of 200 ng/µl) was incubated with N-terminal GST-tagged human SGK3 (final concentration of 10 ng/µl, SignalChem) in reaction buffer (80 mM β-glycerophosphate, 20 mM EGTA, 1 mM MOPS pH 7.0, 1 mM DTT, 1 mM ATP, 5 mM MgCl_2_, and 0.3% NP-40) at 30°C for 50 min. The reaction was stopped by adding LSB followed by heating at 95°C for 5 min. Samples were analyzed by immunoblotting with phospho-specific antibodies.

### GST-AS peptide kinase assay

Freeze-thawed oocytes in ASW (5 oocytes/µl of ASW) were lysed by adding 6.5 volumes of lysis buffer and then incubating samples on ice for 30 min, followed by gentle vortexing. Lysates were centrifuged at 20 k × g at 4°C for 15 min. The supernatant was used for the kinase assay. Mock, Akt-, and SGK-depleted extracts were prepared as described above. For the kinase assay, a reaction mixture containing the GST-AS peptide as a substrate (80 mM β-glycerophosphate, 15 mM MgCl_2_, 20 mM EGTA, 10 mM MOPS, pH 7.0, 1 mM DTT, 1× cOmplete EDTA-free, 2 mM ATP, 0.5 mM sodium orthovanadate, 1 µM okadaic acid, 0.1 mg/ml GST-AS peptide, and 0.3% NP-40) was added to an equal volume of the extract. After incubation at 30°C, the reaction solution was mixed with LSB, heated at 95°C for 5 min, and then analyzed by immunoblotting.

### Statistical analysis

Statistical significance was calculated using the unpaired one-tailed Student’s t-test in Microsoft Excel with StatPlus (AnalystSoft). Statistical significance was defined as p < 0.05.

### Accession numbers

Accession numbers are as follows: LC430700 for sfSGK, AB076395 for sfCdc25, AB060280 for sfMyt1, AB060291 for sfAkt, LC017891 for sfGβ, LC017892 for sfGγ, and LC017893 for the sfPI3K catalytic subunit p110β.

## Data availability

Data supporting the findings of this study are available within the article or upon reasonable requests to the corresponding authors.

## Acknowledgements

We thank E. Okumura for helpful discussions, suggestions, and providing recombinant Cdc25 and Myt1, as well as M. Terasaki for providing the pSP64-S vector. This work was supported by grants-in-aid to T.K. from JSPS [grant numbers 25291043, 16H04783] and Takeda Science Foundation.

## Author contributions

D.H. designed and performed all the experiments. E.H. and K.C. generated the anti-sfSGK-C antibody. All authors discussed the results. D.H. and T.K. wrote the manuscript. T.K. obtained funding and supervised the study.

## Competing interests

The authors declare no competing interests.

## Materials and correspondence

Correspondence and requests for materials should be addressed to D.H. and T.K.

